# Molecular Elucidation of Pancreatic Elastase Inhibition by Baicalein

**DOI:** 10.1101/2020.11.06.371534

**Authors:** Debajeet Ghosh, Sneha Bansode, Rakesh Joshi, Baban Kolte, Rajesh Gacche

**Author notes:** **Corresponding authors**, Prof. Rajesh Gacche, Department of Biotechnology, Savitribai Phule Pune University, Pune, India, Dr. Sneha Bansode, Biochemical Sciences Division, CSIR-National Chemical Laboratory, Pune, India.

## Abstract

The serine protease, elastase exists in various forms and plays diverse roles in the human body. Pharmacological inhibition of elastase has been investigated for its therapeutic role in managing conditions such as diabetes, pneumonia and arthritis. Sivelestat, a synthetic molecule, is the only elastase inhibitor to have been approved by any major drug regulatory authority (PMDA, in this case) – but still has failed to attain widespread clinical usage owing to its high price, cumbersome administration and obscure long-term safety profile. In order to find a relatively better-suited alternative, screening was conducted using plant flavonoids, which yielded Baicalein – a molecule that showed robust inhibition against Pancreatic Elastase inhibition (IC_50_: 3.53 μM). Other than having an IC_50_ almost 1/5^th^ of that of sivelestat’s, baicalein is also cheaper, safer and easier to administer. While Microscale thermophoresis validated baicalein-elastase interaction, enzyme-kinetic studies, molecular docking and molecular dynamic simulation revealed the mode of inhibition to be non-competitive. Baicalein exhibited binding to a distinct allosteric site on the enzyme. The current study demonstrates the elastase inhibition properties of baicalein in an *in-vitro* and *in-silico* environment.

## 1. INTRODUCTION

Elastase is a physiologically relevant serine protease with diverse functions across multiple organ systems. Most notably, its primary function is degrading elastin, collagen and other ECM proteins in arteries, ligaments and lung tissue. The different functions of the enzyme typified its several isotypes – while Neutrophil elastase modulates inflammation and regulates the immune system [1], Pancreatic elastase is a crucial digestive enzyme secreted into the digestive tract.

However, the dysregulation of elastase levels in the blood and certain microenvironments have been proven to be an aggravator behind certain conditions such as Arthritis [2] Pneumonia [3]. The suggested reason behind this might be the imbalance between elastase and its endogenous inhibitors, such as SerpinB1 [5]. Over the years, there has been an exorbitant accumulation of proof that the inhibition of elastase’s several isotypes can be of therapeutic use in conditions which involve such an imbalance [3,4]. Inhibition of neutrophil elastase (NE) has shown promise in the prognosis of several diseases by reducing inflammation-associated tissue damage and reducing immune subversion [1,3]. Having been in the limelight of several decades of research, neutrophil elastase inhibitors have reached the pharmaceutical markets of some countries such as Japan [1,3]. Pancreatic elastase (PE) is mostly known for its use as a gold-standard diagnostic stool test for measuring pancreatic exocrine function. It was only recently reported that like NE, inhibition of pancreatic elastase could also have a beneficial role. It was shown to promote the proliferation of beta islet cells, providing a possible therapy for diabetes [5].

The synthetic drug, sivelestat has a monopoly in the market of elastase inhibition being the only PMDA-approved elastase inhibitor in the market at the moment [6, 15]. The drug molecule has been reported to inhibit both Neutrophil as well as Pancreatic elastase [5]. The inhibition mechanism of sivelestat has been reported to be competitive [7] – it binds to the active site and subsequently prevents enzymatic action on target substrates i.e. ECM proteins. The drug is expensive and requires intravenous administration due to its low GI adsorption. It has quite a few side-effects and the knowledge of its long-term usage safety-profile is limited [8]. These are major hindrances for the Diabetes disease model, which might require patients to regularly use the drug for extensive periods of time. Hence arises the need for an elastase inhibitor which is cheaper, safer and easier to administer.

To find an inhibitor with the aforementioned desirable qualities, we screened plant flavonoids for the property of elastase inhibition. The *in-vitro* inhibitor screening process yielded a molecule, whose binding properties were further studied using Microscale Thermophoresis (MST) and inhibition kinetic(s) assays. To corroborate the *in-vitro* results, *in-silico* measures like Molecular Docking and Molecular Dynamic Simulation were also utilised. As the desired output of this study, we hope to propose the flavonoid molecule – Baicalein – as a potential pancreatic elastase inhibitor.

## 2. MATERIALS & METHODS

### 2.1 Enzyme Screening and Testing

The Michaelis Menten Constant (Km) of Porcine pancreatic elastase (Sigmaaldrich, St. Louis, Missouri, USA) was found out using a suitable chromogenic substrate (Succinyl Ala-Ala-Ala p-nitroanilide, Sigmaaldrich, ≥98% HPLC purity), which yielded a coloured product on enzymatic cleavage. The reaction was set up in 0.5 ml systems containing 0.1M tris buffer (pH 8.0). The Enzyme concentration was 0.001 mg/ml in the system; whereas the Substrate concentration(s) ranged from 0.1 mM to 1.5 mM. The reaction mixture after enzyme, buffer and substrate addition were incubated at 37°C for an hour. Duplicates (200 μL each) were pipetted onto a 96-well plate and OD measured at 405 nm. The molecules tested for elastase inhibition were sourced from Sigmaaldrich – Baicalein (IUPAC: 5,6,7-trihydroxy-2-phenylchromen-4-one) (≥ 98%), Hesperetin (IUPAC: (2*S*)-5,7-dihydroxy-2-(3-hydroxy-4-methoxyphenyl)-2,3-dihydrochromen-4-one) (≥ 95%), 7-Hydroxyflavone (IUPAC: 7-hydroxy-2-phenylchromen-4-one) (≥ 98%), Naringenin (IUPAC: 5,7-dihydroxy-2-(4-hydroxyphenyl)-2,3-dihydrochromen-4-one) (≥ 98%), Galangin (IUPAC: 3,5,7-trihydroxy-2-phenylchromen-4-one) (≥ 95%) and Sivelestat sodium salt hydrate (IUPAC: sodium;2-[[2-[[4-(2,2-dimethylpropanoyloxy)phenyl]sulfonylamino]benzoyl]amino]acetate) (≥98%). The previously mentioned 0.5 mL system was used for testing their inhibitory capacity. The working concentrations of the flavonoids ranged from 1 μM to 125 μM (depending on the extent of inhibition). The Substrate and enzyme concentrations were 0.2 mM and 0.001 mg/ml respectively. It was expected that with increasing inhibition, product formation and the related O.D. will decrease. The IC_50_s for baicalein and sivelestat were calculated by using a best-fit non-linear regression curve in Graph Pad Prism [32]. The same *in-vitro* system was utilized for producing data for the triple-intersection Lineweaver-Burk Plot.

### 2.2 MicroScale Thermophoresis

Fluorescent labelling of the enzyme was done before proceeding with Microscale Thermophoresis. The protein was tagged with NT-647-NHS fluorescent dye using the Monolith NT™ Protein Labelling Kit. Microscale thermophoresis was carried out using the Monolith NT 115 instrument from Nanotemper technologies [33]. Labelled pancreatic elastase enzyme was kept constant at a concentration of (20 μM) whereas the concentration of unlabelled partner baicalein was (0.0153 to 125 μM) in 1× PBS. Using a capillary set (Nanotemper technologies), the samples were loaded one by one into the machine, ensuring the presence of the sample in the mid-section of each capillary. Binding and Ki was detected by proceeding with microscale thermophoresis. LED power of 100% and MST power of 60% were used for this experiment. Data analysis was carried out using NTanalysis (Nanotemper technologies, Munchen, Germany).

### 2.3 Molecular Docking, Visualisation and ADMET testing

For Molecular docking, Autodock v4.2 [18], PatchDock [17] and Autodock Vina [16] were used. The enzyme (PDB ID: 3EST) and inhibitor SDF (PubChem ID: 5281605; FlavoDb Accession Number: FD014805 [40]) files were obtained and processed for Molecular Docking using Autodock Tools v4.2 via PyRx v0.8 [36]. Autogrid was set to ‘maximized’ for Blind Docking; for active site docking the grid size for was set to cover the active site residues. Prior to this, the active site residues were labelled using PyRx itself. Blind Docking was done using Autodock and Autodock Vina, whereas Geometric-Score based Patch docking was done using PatchDock online server. The obtained docked poses and intermolecular interactions were gathered and analysed using Pymol (PyMol Molecular Graphics System, version 1.2r3 pre, Schrodinger LLC). To perform ADME and predictive toxicity testing, Swiss-ADME tool [21] and Pro-Tox II tool [22] were used respectively. To find interacting residues Pymol, Plexview [20] and Discovery Studio (v19.1) [35] were used. LigPlot+ [39] was also utilised for visualisation.

### 2.4 Molecular Dynamic Simulation

The first pose obtained from Molecular Docking (which corresponded to allosteric site 1) was utilised in Molecular Dynamic Simulation. Ligand Parameterization was done, after adding Hydrogen atoms (Discovery Studio v19.1), by the LigParGen online server [36]. NAMD v2.14b2 [37] was used to conduct the MD simulation in a CHARMM forcefield. VMD v1.9.3 [38] was used for visualisation. The ‘Add Solvation Box’ plugin in VMD was used to create an environment with TIP3P water molecules (Jorgensen, Chandrasekhar, Madura, Impey, Klein, 1983) with a Box Padding of 10 Å in all the 3 axes. Followed by this, the ‘Add ions’ plugin was utilised to neutralize the free charges in the system with the ‘Autoionize’ option. First, minimization was conducted for 10,000 steps followed by equilibration for 50,000 steps in an NPT ensemble (utilising Langevin Piston and Langevin temperature control). The Production run was done using an NPT ensemble for 25 ns for the protein-ligand complex as well as the protein alone (timestep = 2 fs). Minimization, equilibration and production run timespans were the same for both Protein-ligand complex and only protein. Periodic Boundary Functions were utilised and Particle Mesh-Ewald Electrostatics was used to calculate long-range electrostatics. The plots were made using the NAMDPlot plugin in VMD. The superimposition images were made using PyMol.

### 2.5 Statistical Analysis

All Statistical analysis and Plotting were done using GraphPad Prism v6.0 [32]. Error bars represent S.D. and P-value less than 0.05 was considered significant. For statistical significance, a two-tailed unpaired T-test was conducted. All *in-vitro* experiments were conducted in triplicates.

## 3. RESULTS

### 3.1 Baicalein exhibited better Pancreatic Elastase inhibition than Sivelestat

A few selected plant flavonoids, previously reported to possess the properties of anti-inflammation and broad-spectrum enzyme inhibition [9 to 13], were tested in an *in-vitro* system to find a potent Pancreatic Elastase (PE) inhibitor. This system yielded a K_m_ for PE which was comparable to the documented one (1.02 mM and 1.15 mM respectively) [14]. Hence, the chosen system was deemed suitable for conducting further experimentation for the study. The flavonoids tested were Baicalein, Naringenin, Hesperetin, 7-Hydroxyflavone and Galangin. Baicalein showed superior inhibition relative to the rest with an IC_50_ of 3.53 μM (Fig 1). Even at high concentrations, hesperetin (125 μM) and galangin (100 μM) showed maximal inhibition(s) of 31.49% and 23.56% respectively; less than 50%. Naringenin and 7-Hydroxyflavone displayed a non-significant inhibition profile across the range of concentrations that were used for screening. Considering the flavonoids (other than baicalein) failed to produce an IC_50_ at even relatively higher concentrations, the study was conducted further only with baicalein.

**Fig 1.**
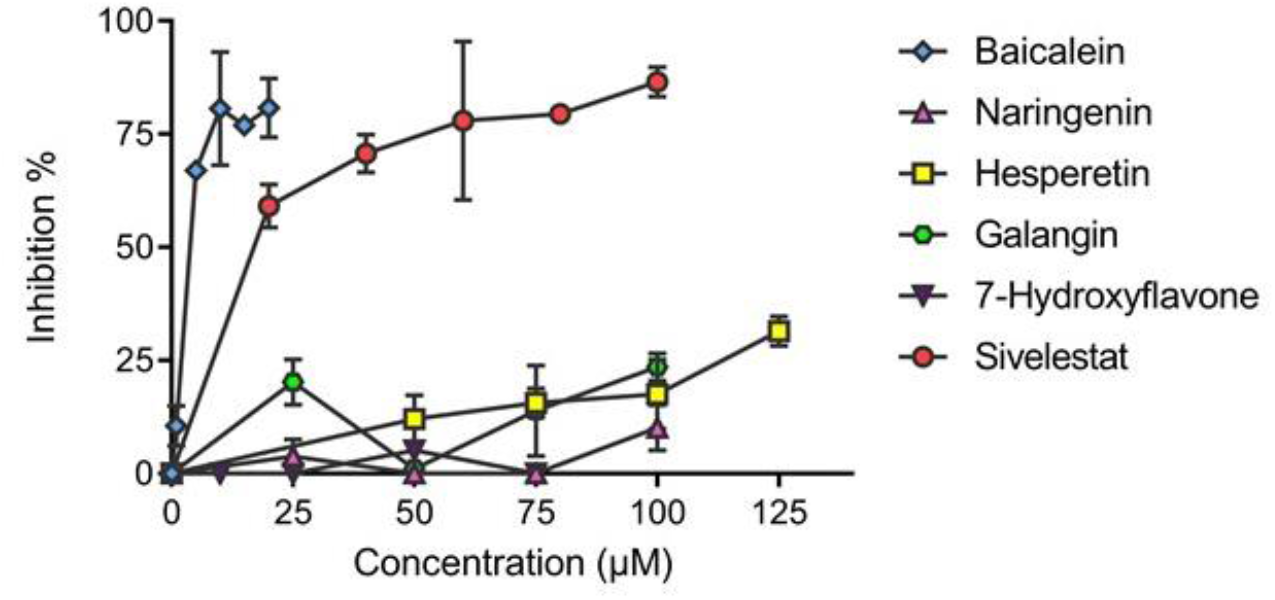
Baicalein displayed better inhibition against elastase than sivelestat as well as other flavonoid inhibitors: Baicalein showed drastically higher inhibition than the other test flavonoids (exhibiting 80.59% inhibition at just 10 μM). The second-best inhibitor was Hesperetin which showed 31.49% at 125 μM. The rest of the inhibitors showed inadequate elastase inhibition even at relatively high concentrations. Sivelestat shows 80% inhibition at around 80-100 μM range, whereas baicalein does it at a mere 10 μM; making it almost an 80-100% better inhibitor when showing maximal inhibition. The IC_50_ value(s) of baicalein and sivelestat were estimated to be 3.53 μM and 15.75 μM, respectively.

Sivelestat is considered as the standard drug inhibitor when performing an elastase inhibition assay. It has been in-use repeatedly as the go-to-standard when performing *in-vitro* and *in-vivo* studies related to elastase inhibition and outcomes [5]. Hence, the molecule’s efficacy to inhibit elastase activity was tested to compare to baicalein (Fig 1). Sivelestat showed significant elastase inhibition at higher concentrations unlike the previously discussed flavonoids (except baicalein). The IC_50_ of sivelestat was estimated to be 15.75 μM. It was interesting to see that baicalein (IC_50_: 3.53 μM) showed around 4.5 times better inhibition than sivelestat. Also, sivelestat required a minimum concentration of around 80 μM to show ~80% inhibition, whereas baicalein did it at a mere 10 μM; making it almost an 8-10X better inhibitor in terms of maximal inhibition.

### 3.2 Microscale thermophoresis validated the binding of Baicalein to Pancreatic Elastase

Microscale thermophoresis was conducted using baicalein (ligand) and fluorescent-tagged PE (target macromolecule) to investigate the binding of baicalein with PE. The increase in baicalein concentration(s) lead to amplified binding of baicalein to elastase; the subsequent binding-affiliated fluorescence quenching can account for the characteristic dip in fluorescent intensity of PE (Fig 2a). Hence, the data achieved from MST was commensurate with our initial observation of baicalein binding to PE. The Ki value was found to be 26.16 nM under tested conditions; amplitude – 39 units and S:N – 8.39.

**Fig 2.**
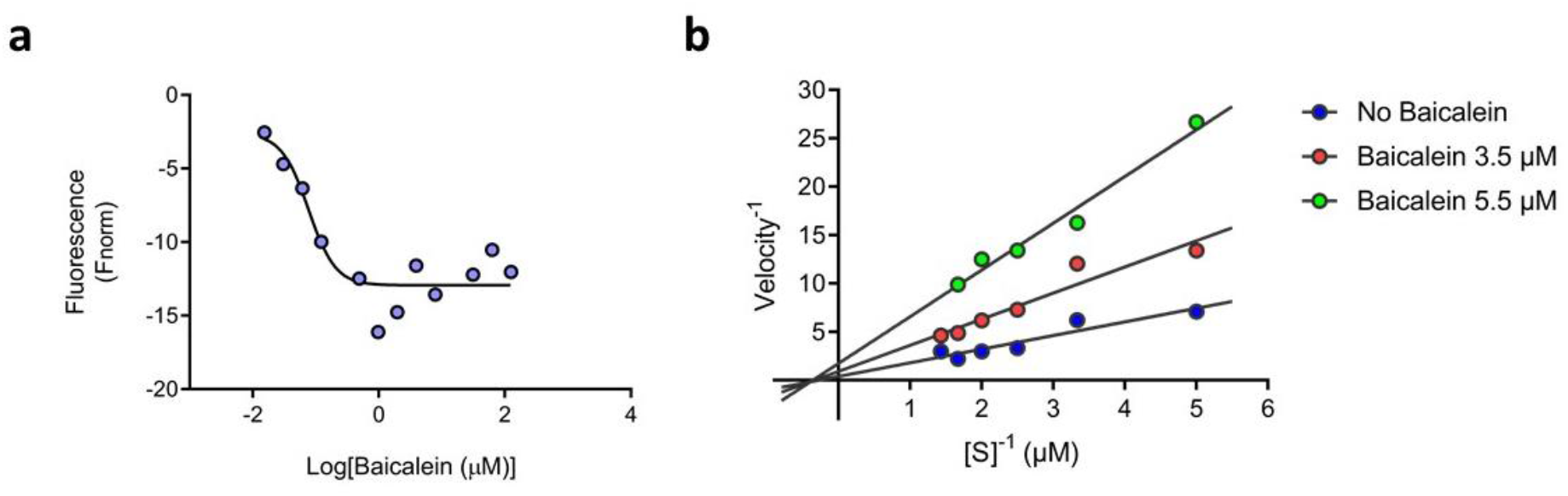
Baicalein binds to Pancreatic Elastase and exhibits non-competitive Inhibition: (a) Microscale Thermophoresis data validated the binding of Baicalein to Pancreatic Elastase– The dip in fluorescence intensity on increasing ligand concentration can be attributed to binding-affiliated quenching. (b) The intersection of the three reciprocal plot lines showed inhibition to be that of noncompetitive fashion; there was hardly any change in the K_m_ on increasing inhibitor concentration.

### 3.3 Baicalein’s mode of inhibition against Pancreatic elastase is non-competitive by nature

Microscale Thermophoresis not only validated the binding of baicalein to Pancreatic elastase, but also eliminated the possibility of uncompetitive inhibition, as the ligand showed binding to the enzyme alone and not the enzyme-substrate complex (a crucial characteristic of uncompetitive inhibition). To test whether the mode of inhibition of baicalein was competitive or non-competitive, a triple intersection Lineweaver-Burk plot was constructed (Fig 2b). The mode of inhibition of baicalein was found to be non-competitive i.e. baicalein binds to an allosteric site of Pancreatic elastase. There was minimal change in K_m_ (if any) on increasing inhibitor concentration – causing us to speculate that baicalein binding does not affect the binding of substrate to enzyme, but affects catalysis instead.

### 3.4 Molecular Docking revealed baicalein’s preference for an allosteric site in Pancreatic elastase

Molecular Docking Studies revealed the possibility of baicalein binding to not only the active site, but also to two other allosteric sites (Fig 3a) – referred to as allosteric site 1 and 2. The average binding energy for allosteric site 1 was found to be lower than that for the active site; −6.825 vs −6.45 kcal/mol (Table 1). Consequently, binding at the allosteric site 1 was thermodynamically favoured and deemed more likely. Blind docking was conducted using Autodock Vina; similar results were found using Autodock and PatchDock Server too (Table S1). Moreover, it was consistently found that the first pose (i.e. the most energetically favourable pose) across the results from different software, were exclusively associated with allosteric site 1. Since blind docking also showed the probable (although less likely) binding of baicalein to the active site, active-site docking was conducted – with baicalein and also sivelestat (for comparison). As sivelestat is known to be a competitive inhibitor, its affinity for the active site should be more than baicalein’s. Active site docking results (Fig 3b) proved that sivelestat indeed had better affinity towards the active site when compared to baicalein (−7.1 kcal/mol vs −6.5 kcal/mol).

**Table 1.** Blind Docking Results (Autodock Vina)

| Site | Pose(s) | Interacting residues | Average Energy (kcal/mol) |
| --- | --- | --- | --- |
| Active Site | 6, 8 | His57, Ser195, Gly193, Val216, Gln192 | -6.450 |
| Allosteric Site 1 | 1, 2, 4, 5 | His200, Leu120, Ser29, Val122, Tyr207, Tyr137 | -6.825 |
| Allosteric Site 2 | 3, 9 | Asn74, Gln34, Tyr82, Asp65 | -6.700 |

**Fig 3.**
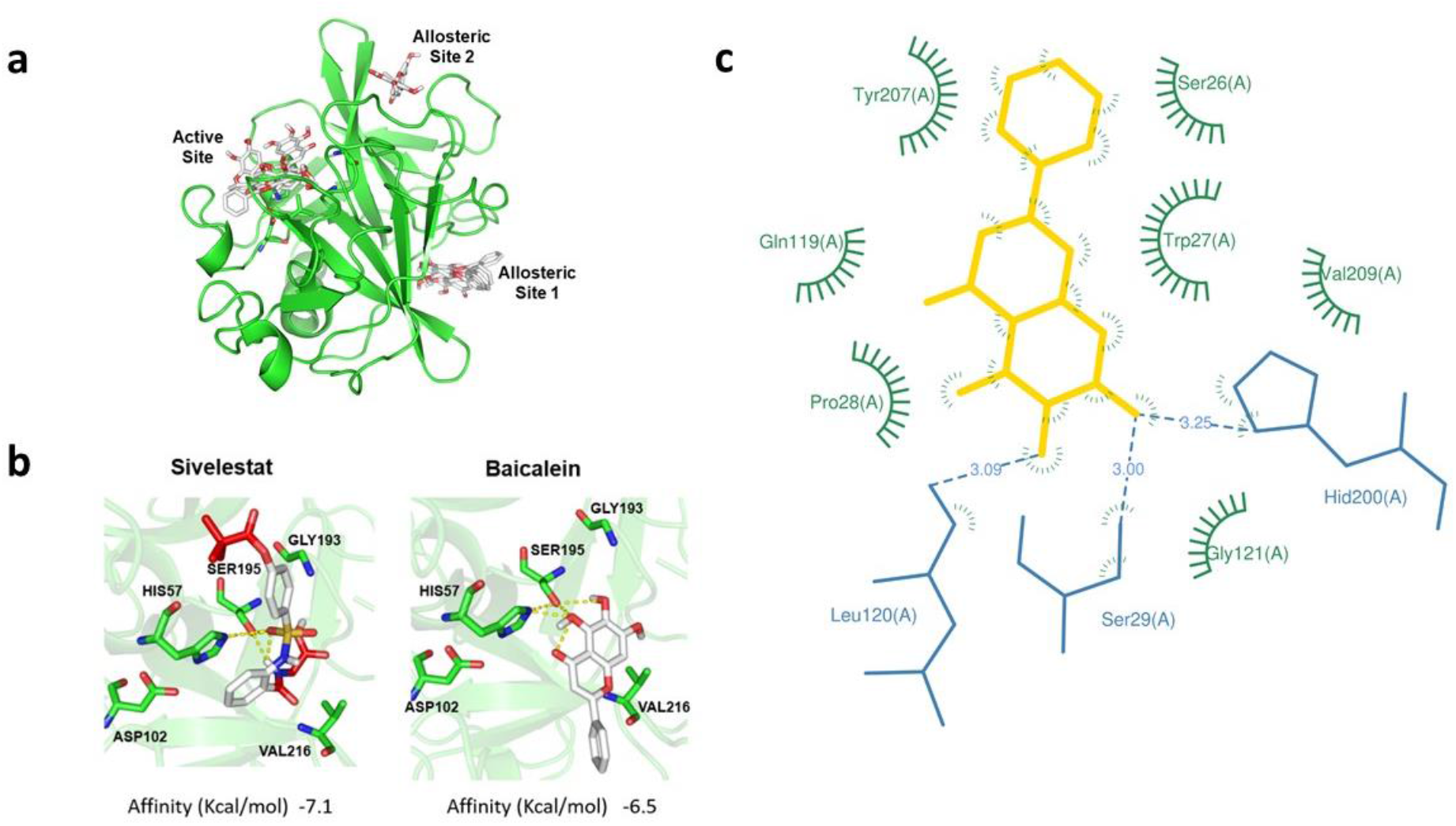
Molecular Docking provided additional proof(s) of allosteric inhibition of Baicalein: (a) Molecular Docking revealed the possibility of Baicalein binding to the active site as well as two other allosteric sites. (b) Active Site docking revealed Sivelestat (left) to have more affinity for the active site than Baicalein (right); −7.1 vs −6.5 kcal/mol. (C) LigPlot+ was used to predict the interacting residues from the Docking results that were obtained (Table 1). Blue lines represent H-bonding; green circular spokes represent Hydrophobic interaction.

The interacting residues of allosteric site 1 (suspected to be the site of baicalein binding) were found out across all the obtained poses (Table 1) and the residue-interactome for the first pose was summed up pictorially (Fig 3c). Out of the various interacting residues, the most conspicuously and commonly found residues across the different poses were Ser29 and Tyr207. Plexview Online software [22] was used to verify the interaction, and it corroborated the residue interactome data. It showed Tyr207 to be involved in pi-pi bonding with the carbon atoms of baicalein, whereas Ser29 interacted via H-bonds (Fig 3c).

### 3.5 Molecular Dynamic Simulation corroborated the allosteric binding of baicalein to Pancreatic elastase

To elucidate the interaction between baicalein and PE at an atomic resolution, MD simulation studies were conducted using the PE-baicalein complex (baicalein at allosteric site 1). The RMSD of the Protein backbone in the complex (PE + baicalein) showed a pronounced increase in RMSD over the non-bound Protein (PE only) (Fig 4A). The increase in the mean RMSD of the PE-baicalein complex vs PE only (1.92 Å vs 1.64 Å) was also statistically significant; p-value <0.0001. This can be attributed to ligand-induced changes in the structure of Pancreatic elastase, which is only possible if the ligand binds to the protein. A second run of simulation with PE was conducted to eliminate the possibility of the difference in RMSD (between the curves) being a false-positive result. However, it was found that the RMSD curve for PE (unbound protein) across multiple runs did not significantly deviate from the displayed graph (Fig 4a) unless baicalein was introduced to the *in-silico* environment. The overall RMSF curve was also higher for PE-baicalein complex, with the largest differences being for residues Asn221 and Ser217 (Fig 4b). The Total Energy Curves for the bound and unbound protein was found out using NAMDPlot (Fig 4c).

**Fig 4.**
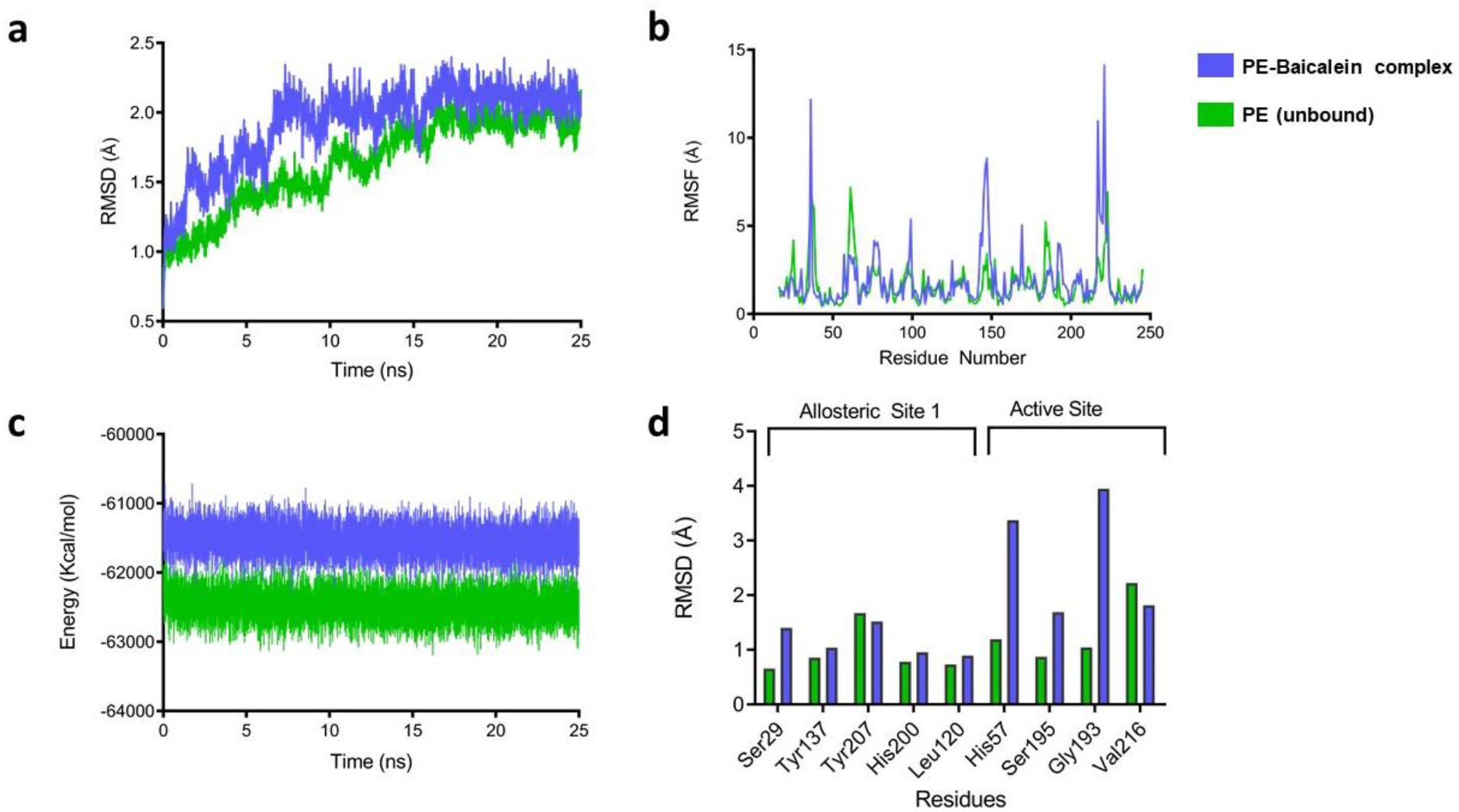
Molecular Docking studies elucidated interaction between Pancreatic elastase and baicalein at an atomic resolution: (a) Pancreatic elastase (PE) – baicalein complex (blue) showed a pronounced increase in the RMSD curve compared to unbound PE (green). The mean RMSD of the complex’s protein backbone was significantly higher than that of the unbound protein – p < 0.0001. (b) The protein-ligand complex’s RMSF curve was also conspicuously higher than the unbound protein. (c) The Total Energy Curve(s) of the bound and unbound protein during the course of the simulation. (d) RMSD of Individual residues present in Allosteric Site 1 and Active Site – unbound vs protein-ligand complex.

The RMSD curve of baicalein revealed a sharp rise starting around 8 ns which was consistent for the rest of the simulation (Fig S2a). This spike in RMSD reveals there to be a significant change in the position/physical properties of the ligand – an occurrence expected when a ligand undergoes positional adjustments to fit into the site of a larger macromolecule. Hence it can be speculated that the event of baicalein binding is occurring at 8 ns. The phase between 4.5 ns and 6.3 ns, where RMSD rises for a brief period of time can be hypothesised to be an ‘adjustment phase’ per se – where baicalein temporarily interacts with the residues of Allosteric Site 1 before finally binding at 8 ns. A similar shift in the ligand RMSD curve was also noticed when MD simulation was conducted with the complex in an Amber force field (by the online server PlayMolecule.org), but the event of RMSD spike (or ligand-target binding) was suspected to be at 19-20 ns instead (Fig S2b).

Although baicalein was not seen to bind to the active site, baicalein was previously shown to affect the active site through allosterism. Hence, RMSD of Allosteric Site 1 as well as Active Site residues were measured (Fig 4d). The increase in RMSD in the residues of the protein-ligand complex was especially high in the active-site residues His57 and Gly193. Thus, it can be deemed that baicalein’s binding to the Allosteric Site 1 has a discernible impact on the physical behaviour of the catalytic active site residues. This is to be expected if baicalein indeed has an impact on the enzymatic activity of the target enzyme PE, which it does via allosterism. On inspecting the movement of baicalein inside allosteric site 1, it was seen to move around inside the site throughout simulation – consistent interaction between any one residue and baicalein could not be visualised. Rather, baicalein seemed to cycle across all the allosteric site 1 residues (Table 1) via transient interactions. It assumed structures and coordinates similar to the poses 1,2,4 and 5 (Table 1). It was consistently present inside Allosteric site 1 for the entire simulation and interacted with Tyr207 the most out of all the other residues. The before and after results of MD simulation showed a relative shift in the position of baicalein, its interacting residues (Fig 5b) and the overall structure of the protein (Fig 5a).

**Fig 5.**
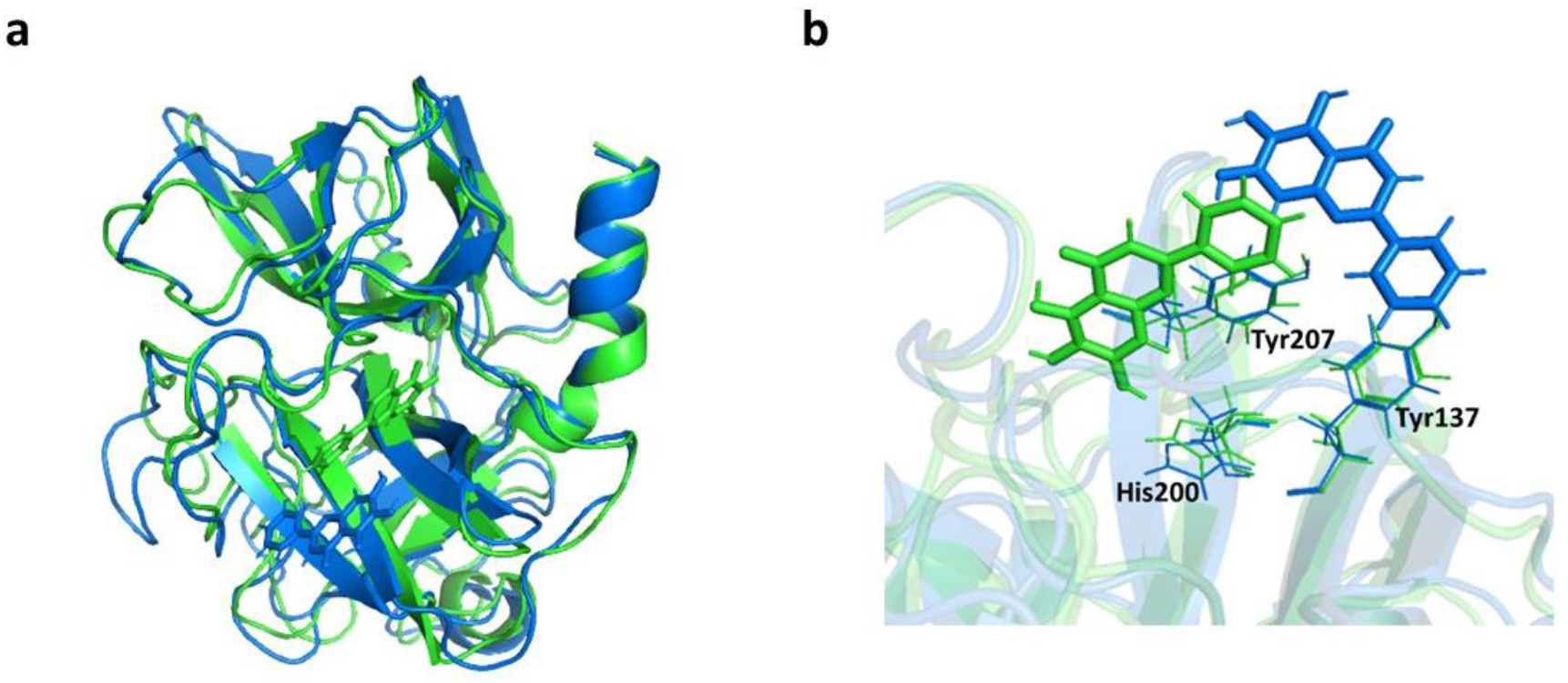
Pre-MD and Post-MD visualisation of protein and ligand: (a) A superimposition of the pre-MD (Green) and post-MD (Blue) images of PE −baicalein complex. The cartoon represents the secondary structure of the protein chain; the licorice-represented molecules correspond to baicalein. (b) The change in position of baicalein before and after MD simulation with respect to three crucial allosteric site 1 residues.

### 3.6 ADMET analysis showed baicalein to have high drug-likeness and low toxicity

To assess the drug-like qualities of baicalein – ADME analysis (SwissADME [21]) was done using sivelestat for comparison (Table 2), which is a PMDA-approved elastase inhibitor (awaiting FDA approval, [6]). It was found that baicalein followed all of Lipinski’s rules (0 violations) and the bioavailability score of baicalein was comparable to sivelestat. However, baicalein was predicted to have high GI absorption, which was low for sivelestat. Predictive toxicity testing, with ProTox-II [22], illustrated baicalein’s toxicity profile to be low and comparable to sivelestat. Hence, it was conclusively seen that baicalein met all the pre-requisites required by a molecule to quality it as a drug; it might even be a more suitable candidate than sivelestat in this aspect, considering its higher rate of GI absorption. Corroborating the predictive *in silico* results, previous reports have suggested that baicalein is well tolerated in patients, is non-toxic and has a favourable safety profile [30].

**Table 2.**
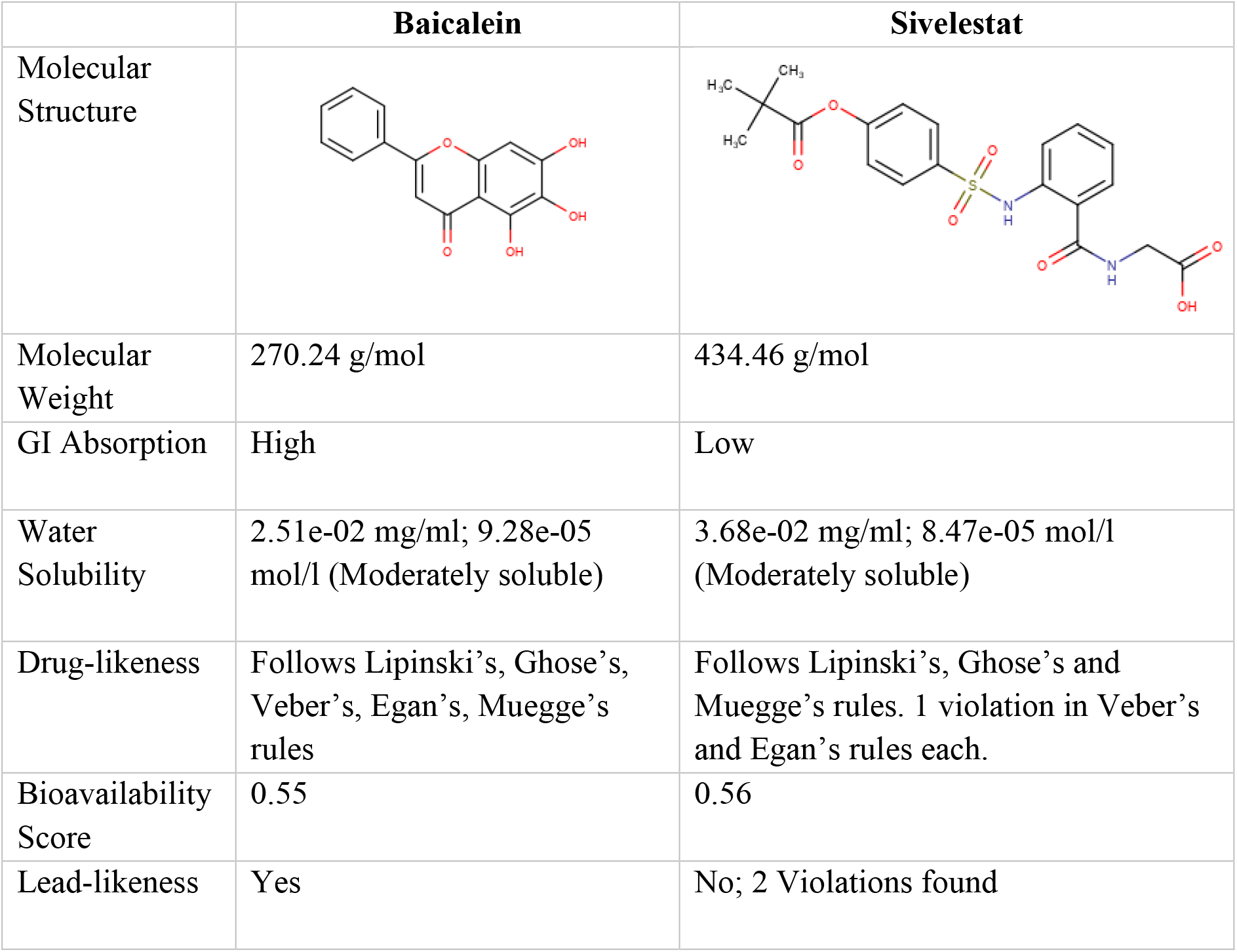
Comparison of ADMET analysis between Baicalein and Sivelestat.

|  | Baicalein | Sivelestat |
| --- | --- | --- |
| Molecular Structure |  |  |
| Molecular Weight | 270.24 g/mol | 434.46 g/mol |
| GI Absorption | High | Low |
| Water Solubility | 2.51e-02 mg/ml; 9.28e-05 mol/l (Moderately soluble) | 3.68e-02 mg/ml; 8.47e-05 mol/l (Moderately soluble) |
| Drug-likeness | Follows Lipinski's, Ghose's, Veber's, Egan's, Muegge's rules | Follows Lipinski's, Ghose's and Muegge's rules. 1 violation in Veber's and Egan's rules each. |
| Bioavailability Score | 0.55 | 0.56 |
| Lead-likeness | Yes | No; 2 Violations found |

## DISCUSSION

Plant flavonoids are preferred as a source of therapy, due to the lower occurrence of side effects and long-term safety concerns. Besides this, they are also well perceived by patients, due to the advertent conceptualising of ‘natural’ being the best. The study initially aimed at finding natural alternatives to sivelestat for Pancreatic elastase inhibition. Inhibition of the protease tends to show islet hyperplasia, a phenomenon which may be utilised for therapeutic purposes in diabetes [5].

Baicalein is a plant flavonoid derived from the roots of the medicinal Chinese herb, *Scutellaria baicalensis*. It has been used historically in Chinese herbal supplements for improving hepatic health. Otherwise, baicalein has also been found to be relevant to several aspects of human health. It has shown to inhibit several inflammatory prostaglandins, exhibiting properties of a promising anti-inflammatory [23]. It also acts as an anxiolytic and estrogen-inhibitor [24]. However, it mostly came to the limelight when its profound multi-faceted action against cancer was discovered [25].

Through this study, we elucidate a novel property of baicalein – elastase inhibition. The molecule showed superior inhibition against Pancreatic elastase, with an IC_50_ of 3.53 μM - better than sivelestat by several folds. The fact that baicalein is a strong elastase inhibitor, may construe several possibilities – Baicalein administration will probably exhibit islet hyperplasia too, just like sivelestat, serpinB1 and other elastase inhibitors [5] allowing the opportunity for it to be a potent therapy in certain conditions such as diabetes. Also, baicalein may be better tolerated than sivelestat, being a plant product. Sivelestat has a considerable number of side effects [8] – which may be ameliorated on shifting from a sivelestat-based therapy to a baicalein one.

As mentioned previously, baicalein has over the past decades seen utility in a wide variety of human conditions for its beneficial attributes towards human health. There are several studies which prove that baicalein’s anti-inflammatory properties help improve beta-cell function, prevent beta-cell damage [26] and combat hyperglycemia [27]. Baicalein is also much cheaper than sivelestat [28, 29] – former being priced almost 1/50^th^ of the latter. All in all, baicalein can be said to be a cheaper and stronger natural alternative to sivelestat, which combats the derogatory effects of Diabetes by not only promoting elastase inhibition but also in other ways – improving beta-cell viability, preventing hyperlipidemia-induced damage to beta cells, promoting GSIS [26], reducing hyperglycemia [29] – none of which sivelestat do. Considering that sivelestat administration is usually intratracheal or intraperitoneal – both of which require medical assistance – its administration makes its usage difficult for Diabetes therapy, where patients are required to be under drug administration on a daily basis. Predictive bioinformatics and pharmacokinetics showed baicalein to have higher GI absorption than sivelestat. Baicalein’s administration is also usually via ingestion (tablet/powder form) as per previous clinical trials [25, 30] – showing baicalein’s administration and bioavailability to be better than sivelestat’s. Assessing the drug-administration is of crucial importance in lifestyle disorders such as diabetes where daily-medication is often administered by the patients themselves. Also, the suggested remedy of gradual beta-cell proliferation would require long-term usage of the drug, making administration all the more important.

Sivelestat is a competitive inhibitor. As per the results obtained from the *in-vitro* and *in-silico* findings, baicalein demonstrates the properties of a non-competitive inhibitor. Allosteric modulation has been previously reported for baicalein [31]. Non-competitive inhibition underlines the possibility of the inhibitor molecule binding to either the enzyme alone or the enzyme-substrate complex. However, MST data elucidated that the inhibitor showed binding to the enzyme alone, and not the Enzyme-Substrate complex. On binding to an allosteric site in the unbound enzyme, it most likely affects the catalysis but not the binding (as there seems to be hardly any change in K_m_, Fig 2B). A hallmark difference between competitive and non-competitive inhibition is that the former is affected by the concentration of the substrate; the latter however is not. For instance, in high levels of elastase accumulation a competitive inhibitor (like sivelestat) may not work as well as a non-competitive inhibitor (like baicalein). Hypothetically, this is yet another ground where baicalein claims superiority over sivelestat. Other than inhibiting Pancreatic elastase, sivelestat also inhibits the immunologically-relevant enzyme – Neutrophil elastase. Considering the similarity in structure between the two elastase isotypes it might be speculated that baicalein probably inhibits neutrophil elastase also.

In conclusion, baicalein’s superior inhibition property might find therapeutic opportunity through inhibition of Pancreatic elastase. The flavonoid has been subjected to extensive clinical trials for testing its other beneficial attributes, and has also shown an impressive safety profile. Other than that, this new found property of elastase inhibition, might explain the physiological basis of side effects that may occur (although rarely) in the case of high-dosage baicalein administration. Baicalein’s pure non-competitive inhibition with limited variance in K_m_ was also unique. Although the results leave little doubt about allosteric interactions(s), the possibility of baicalein binding to both the active site and allosteric site is yet to be explored.

## Acknowledgements

Authors are thankful to Savitribai Phule Pune University (SPPU), Pune for providing the facility and requirements needed for the study. We are also thankful to Dr. Rohan Meshram for his help in the MD simulation study.

## REFERENCES

1. Döring G. (1994). The role of neutrophil elastase in chronic inflammation. American journal of respiratory and critical care medicine, 150(6 Pt 2), S114–S117. https://doi.org/10.1164/ajrccm/150.6_Pt_2.S114

2. Muley, M. M., Reid, A. R., Botz, B., Bölcskei, K., Helyes, Z., & McDougall, J. J. (2016). Neutrophil elastase induces inflammation and pain in mouse knee joints via activation of proteinase-activated receptor-2. British journal of pharmacology, 173(4), 766–777. https://doi.org/10.1111/bph.

3. Domon, H., Nagai, K., Maekawa, T., Oda, M., Yonezawa, D., Takeda, W., Hiyoshi, T., Tamura, H., Yamaguchi, M., Kawabata, S., & Terao, Y. (2018). Neutrophil Elastase Subverts the Immune Response by Cleaving Toll-Like Receptors and Cytokines in Pneumococcal Pneumonia. Frontiers in immunology, 9, 732. https://doi.org/10.3389/fimmu.2018.00732

4. Dharwal, V., & Naura, A. S. (2018). PARP-1 inhibition ameliorates elastase induced lung inflammation and emphysema in mice. Biochemical pharmacology, 150, 24–34. https://doi.org/10.1016/j.bcp.2018.01.027

5. El Ouaamari, A., Dirice, E., Gedeon, N., Hu, J., Zhou, J. Y., Shirakawa, J., Hou, L., Goodman, J., Karampelias, C., Qiang, G., Boucher, J., Martinez, R., Gritsenko, M. A., De Jesus, D. F., Kahraman, S., Bhatt, S., Smith, R. D., Beer, H. D., Jungtrakoon, P., Gong, Y.,… Kulkarni, R. N. (2016). SerpinB1 Promotes Pancreatic β Cell Proliferation. Cell metabolism, 23(1), 194–205. https://doi.org/10.1016/j.cmet.2015.12.001

6. KEGG Drug Database, New Drug Approvals in the USA, Europe and Japan, accessed 10th August 2020: <https://www.genome.jp/kegg/drug/br08328.html?id=D01918>

7. Kawabata, K., Suzuki, M., Sugitani, M., Imaki, K., Toda, M., & Miyamoto, T. (1991). ONO-5046, a novel inhibitor of human neutrophil elastase. Biochemical and biophysical research communications, 177(2), 814–820. https://doi.org/10.1016/0006-291x(91)91862-7

8. Aikawa, N., & Kawasaki, Y. (2014). Clinical utility of the neutrophil elastase inhibitor sivelestat for the treatment of acute respiratory distress syndrome. Therapeutics and clinical risk management, 10, 621–629. https://doi.org/10.2147/TCRM.S65066

9. Cho J. (2006). Antioxidant and neuroprotective effects of hesperidin and its aglycone hesperetin. Archives of pharmacal research, 29(8), 699–706. https://doi.org/10.1007/BF02968255

10. Wolfram, J., Scott, B., Boom, K., Shen, J., Borsoi, C., Suri, K., Grande, R., Fresta, M., Celia, C., Zhao, Y., Shen, H., & Ferrari, M. (2016). Hesperetin Liposomes for Cancer Therapy. Current drug delivery, 13(5), 711–719. https://doi.org/10.2174/1567201812666151027142412

11. Bie, B., Sun, J., Guo, Y., Li, J., Jiang, W., Yang, J., Huang, C., & Li, Z. (2017). Baicalein: A review of its anti-cancer effects and mechanisms in Hepatocellular Carcinoma. Biomedicine & pharmacotherapy = Biomedecine & pharmacotherapie, 93, 1285–1291. https://doi.org/10.1016/j.biopha.2017.07.068

12. Jin, J., Chen, Y., Wang, D., Ma, L., Guo, M., Zhou, C., & Dou, J. (2018). The inhibitory effect of sodium baicalin on oseltamivir-resistant influenza A virus via reduction of neuraminidase activity. Archives of pharmacal research, 41(6), 664–676. https://doi.org/10.1007/s12272-018-1022-6

13. Bao, L., Liu, F., Guo, H. B., Li, Y., Tan, B. B., Zhang, W. X., & Peng, Y. H. (2016). Naringenin inhibits proliferation, migration, and invasion as well as induces apoptosis of gastric cancer SGC7901 cell line by downregulation of AKT pathway. Tumour biology: the journal of the International Society for Oncodevelopmental Biology and Medicine, 37(8), 11365–11374. https://doi.org/10.1007/s13277-016-5013-2

14. Km information; Elastase, pancreatic from porcine pancreas, Sigma-Aldrich, accessed 10th August, 2020. <https://www.sigmaaldrich.com/content/dam/sigma-aldrich/docs/Sigma/Product_Information_Sheet/e0127pis.pdf>

15. Sivelestat launch in Japanese markets: https://www.thepharmaletter.com/article/ono-launches-elastase-inhibitor-in-japan

16. Trott, O., & Olson, A. J. (2010). AutoDock Vina: improving the speed and accuracy of docking with a new scoring function, efficient optimization, and multithreading. Journal of computational chemistry, 31(2), 455–461. https://doi.org/10.1002/jcc.21334

17. Schneidman-Duhovny, D., Inbar, Y., Nussinov, R., & Wolfson, H. J. (2005). PatchDock and SymmDock: servers for rigid and symmetric docking. Nucleic acids research, 33(Web Server issue), W363–W367. https://doi.org/10.1093/nar/gki481

18. Morris, G. M., Huey, R., Lindstrom, W., Sanner, M. F., Belew, R. K., Goodsell, D. S., & Olson, A. J. (2009). AutoDock4 and AutoDockTools4: Automated docking with selective receptor flexibility. Journal of computational chemistry, 30(16), 2785–2791. https://doi.org/10.1002/jcc.21256

19. Phillips, J. C., Braun, R., Wang, W., Gumbart, J., Tajkhorshid, E., Villa, E., Chipot, C., Skeel, R. D., Kalé, L., & Schulten, K. (2005). Scalable molecular dynamics with NAMD. Journal of computational chemistry, 26(16), 1781–1802. https://doi.org/10.1002/jcc.20289

20. Plexview Online Software, accessed 10th August, 2020: <https://playmolecule.org/PlexView/>

21. Daina, A., Michielin, O., & Zoete, V. (2017). SwissADME: a free web tool to evaluate pharmacokinetics, drug-likeness and medicinal chemistry friendliness of small molecules. Scientific reports, 7, 42717. https://doi.org/10.1038/srep42717

22. Banerjee, P., Eckert, A. O., Schrey, A. K., & Preissner, R. (2018). ProTox-II: a webserver for the prediction of toxicity of chemicals. Nucleic acids research, 46(W1), W257–W263. https://doi.org/10.1093/nar/gky318

23. Hsieh, C. J., Hall, K., Ha, T., Li, C., Krishnaswamy, G., & Chi, D. S. (2007). Baicalein inhibits IL-1beta- and TNF-alpha-induced inflammatory cytokine production from human mast cells via regulation of the NF-kappaB pathway. Clinical and molecular allergy: CMA, 5, 5.

24. Stefanie Schwartz (9 January 2008). Psychoactive Herbs in Veterinary Behavior Medicine. John Wiley & Sons. pp. 139–. ISBN 978-0-470-34434-7.

25. Liu, H., Dong, Y., Gao, Y., Du, Z., Wang, Y., Cheng, P., Chen, A., & Huang, H. (2016). The Fascinating Effects of Baicalein on Cancer: A Review. International journal of molecular sciences, 17(10), 1681. https://doi.org/10.3390/ijms17101681

26. Fu, Y., Luo, J., Jia, Z., Zhen, W., Zhou, K., Gilbert, E., & Liu, D. (2014). Baicalein Protects against Type 2 Diabetes via Promoting Islet β-Cell Function in Obese Diabetic Mice. International journal of endocrinology, 2014, 846742. https://doi.org/10.1155/2014/846742

27. Ku, S. K., & Bae, J. S. (2015). Baicalin, baicalein and wogonin inhibits high glucose-induced vascular inflammation in vitro and in vivo. BMB reports, 48(9), 519–524. https://doi.org/10.5483/bmbrep.2015.48.9.017

28. Baicalein (98%), Sigmaaldrich website, Accessed 10th August, 2020:<https://www.sigmaaldrich.com/catalog/search?term=491-67-8&interface=CAS%20No.&N=0&mode=partialmax&lang=en&region=IN&focus=product&gclid=Cj0KCQjw7qn1BRDqARIsAKMbHDZhlhbXfRqUONB6f23fcS0aoOm9qJ_EZ_tbWUzAREsqoXBo340qAFQaAlqkEALw_wcB>

29. Sivelestat Sodium Salt Hydrate (98%), Sigmaaldrich website, Accessed 10th August,2020: https://www.sigmaaldrich.com/catalog/product/sigma/s7198?lang=en&region=IN

30. Clinicaltrials.gov (by NIH); A Randomized, Double-blind, Placebo-controlled, Multicenter and Phase IIa Clinical Trial for the Effectiveness and Safety of Baicalein Tablets in the Treatment of Improve Other Aspects of Healthy Adult With Influenza Fever; CSPC ZhongQi Pharmaceutical Technology Co., Ltd.

31. Han, J., Ji, Y., Youn, K., Lim, G., Lee, J., Kim, D. H., & Jun, M. (2019). Baicalein as a Potential Inhibitor against BACE1 and AChE: Mechanistic Comprehension through In Vitro and Computational Approaches. Nutrients, 11(11), 2694. https://doi.org/10.3390/nu11112694

32. GraphPad Prism version 6.00 for Windows, GraphPad Software, La Jolla California USA, <www.graphpad.com>

33. Jerabek-Willemsen, M., Wienken, C. J., Braun, D., Baaske, P., & Duhr, S. (2011). Molecular interaction studies using microscale thermophoresis. Assay and drug development technologies, 9(4), 342–353. https://doi.org/10.1089/adt.2011.0380

34. Dallakyan, S., & Olson, A. J. (2015). Small-molecule library screening by docking with PyRx. Methods in molecular biology (Clifton, N.J.), 1263, 243–250. https://doi.org/10.1007/978-1-4939-2269-7_19

35. Dassault Systèmes BIOVIA, Discovery Studio Modeling Environment, Release 2017, San Diego: Dassault Systèmes, 2016.

36. Dodda, L. S., Cabeza de Vaca, I., Tirado-Rives, J., & Jorgensen, W. L. (2017). LigParGen web server: an automatic OPLS-AA parameter generator for organic ligands. Nucleic acids research, 45(W1), W331–W336. https://doi.org/10.1093/nar/gkx312.

37. Phillips, J. C., Braun, R., Wang, W., Gumbart, J., Tajkhorshid, E., Villa, E., Chipot, C., Skeel, R. D., Kalé, L., & Schulten, K. (2005). Scalable molecular dynamics with NAMD. Journal of computational chemistry, 26(16), 1781–1802. https://doi.org/10.1002/jcc.20289

38. Humphrey, W., Dalke, A., & Schulten, K. (1996). VMD: visual molecular dynamics. Journal of molecular graphics, 14(1), 33–28. https://doi.org/10.1016/0263-7855(96)00018-5

39. Laskowski, R. A., & Swindells, M. B. (2011). LigPlot+: multiple ligand-protein interaction diagrams for drug discovery. Journal of chemical information and modeling, 51(10), 2778–2786.https://doi.org/10.1021/ci200227u

40. Kolte, B.S., Londhe, S.R., Bagul, K.T. et al. (2019). FlavoDb: a web-based chemical repository of flavonoid compounds. 3 Biotech 9, 431 https://doi.org/10.1007/s13205-019-1962-7

