## Supplementary figure 1, 2, and table for "Molecular Elucidation of Pancreatic Elastase Inhibition by Baicalein"

### Slide 1
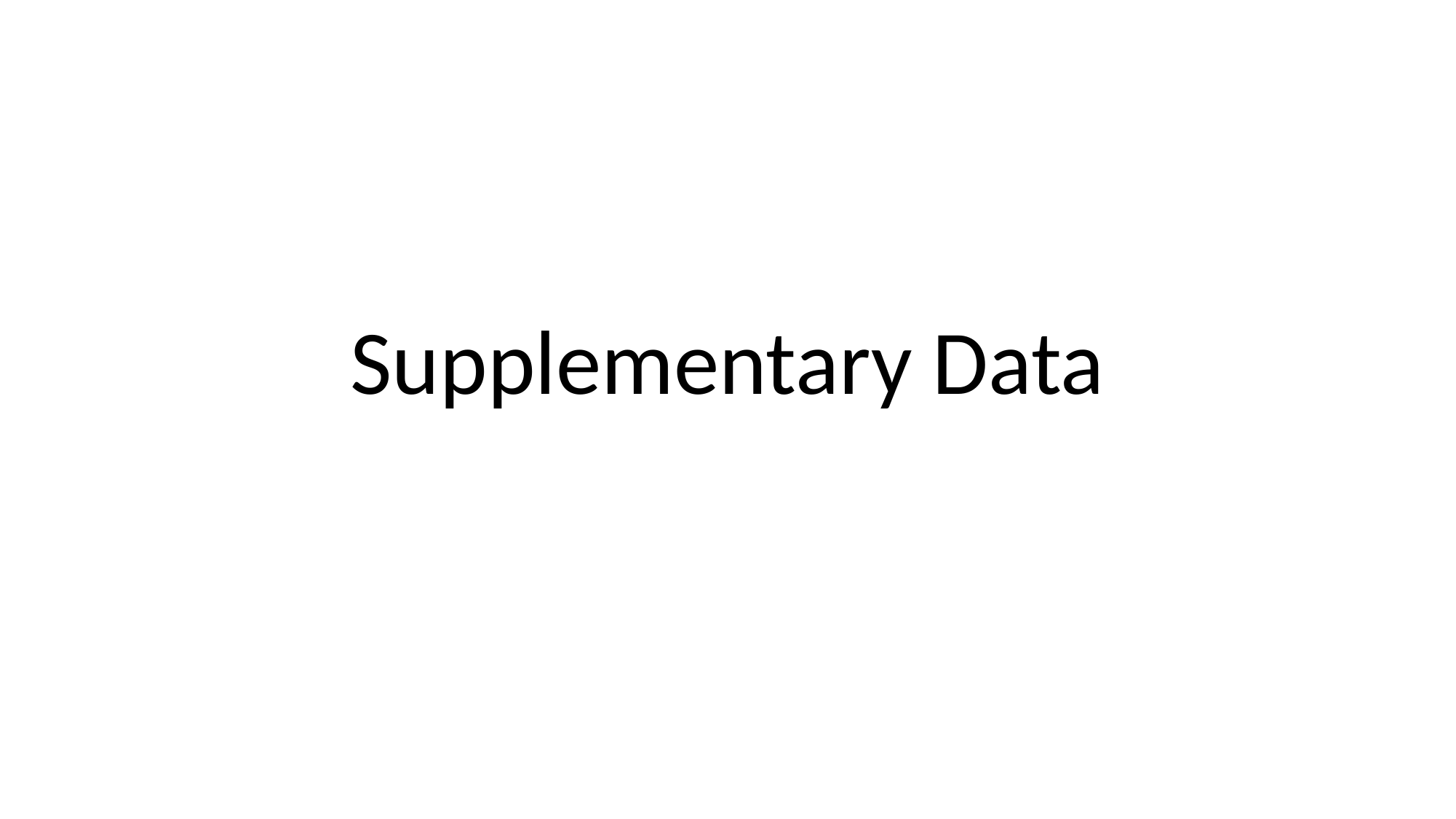

#### Supplementary Data

### Slide 2
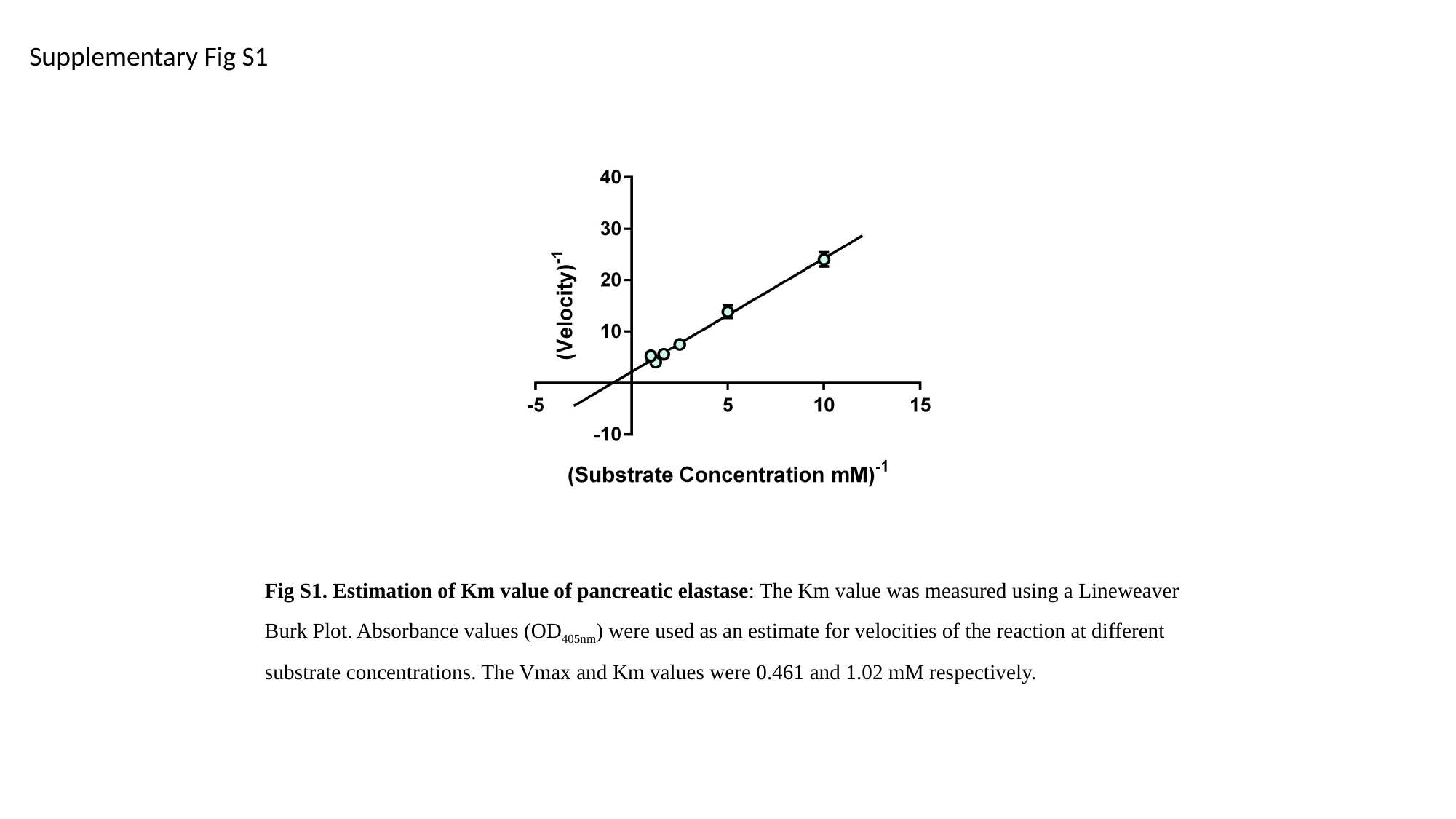

Supplementary Fig S1
Fig S1. Estimation of Km value of pancreatic elastase: The Km value was measured using a Lineweaver Burk Plot. Absorbance values (OD_405nm_) were used as an estimate for velocities of the reaction at different substrate concentrations. The Vmax and Km values were 0.461 and 1.02 mM respectively.

### Slide 3
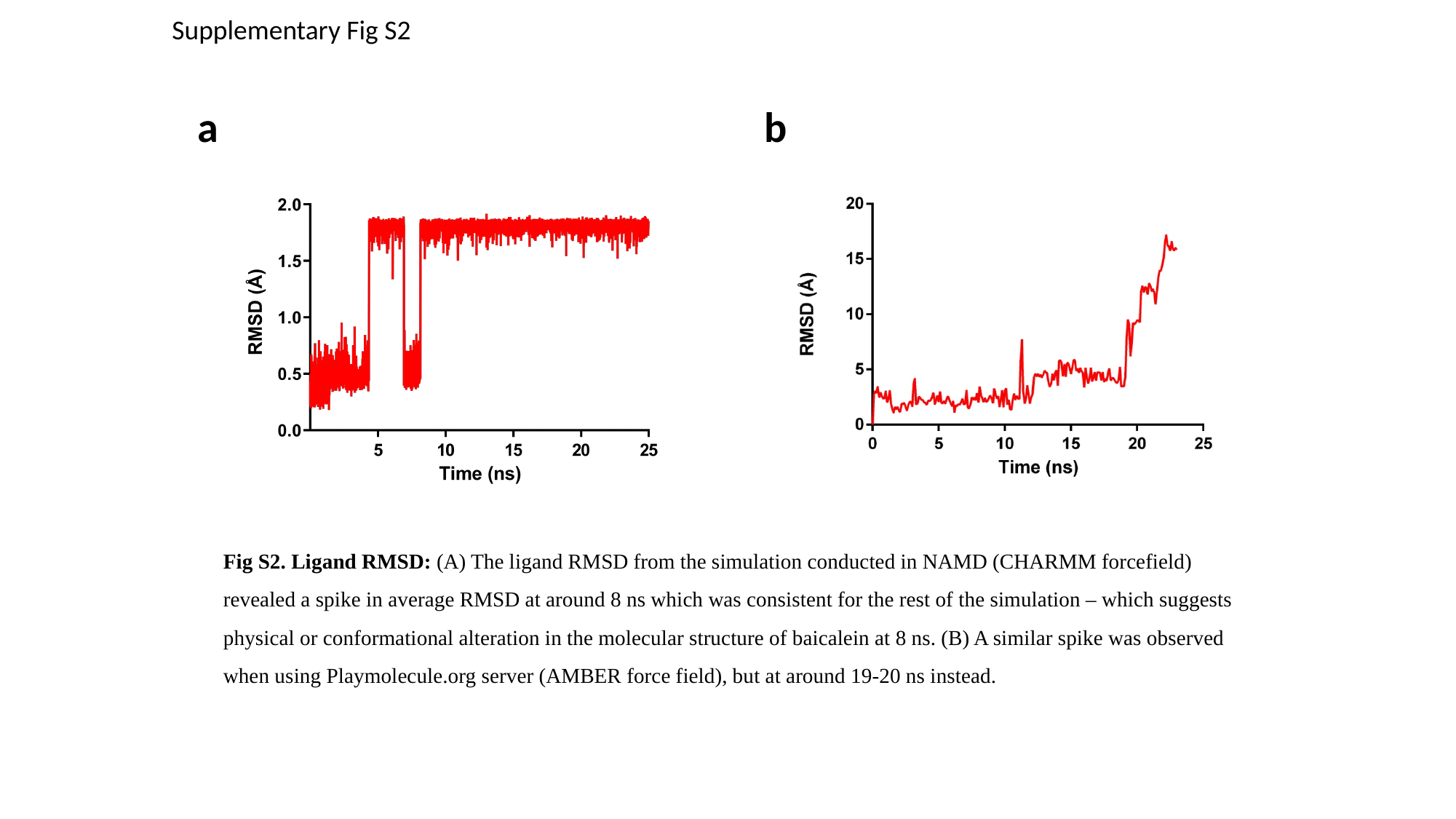

Supplementary Fig S2
a
b
Fig S2. Ligand RMSD: (A) The ligand RMSD from the simulation conducted in NAMD (CHARMM forcefield) revealed a spike in average RMSD at around 8 ns which was consistent for the rest of the simulation – which suggests physical or conformational alteration in the molecular structure of baicalein at 8 ns. (B) A similar spike was observed when using Playmolecule.org server (AMBER force field), but at around 19-20 ns instead.

### Slide 4
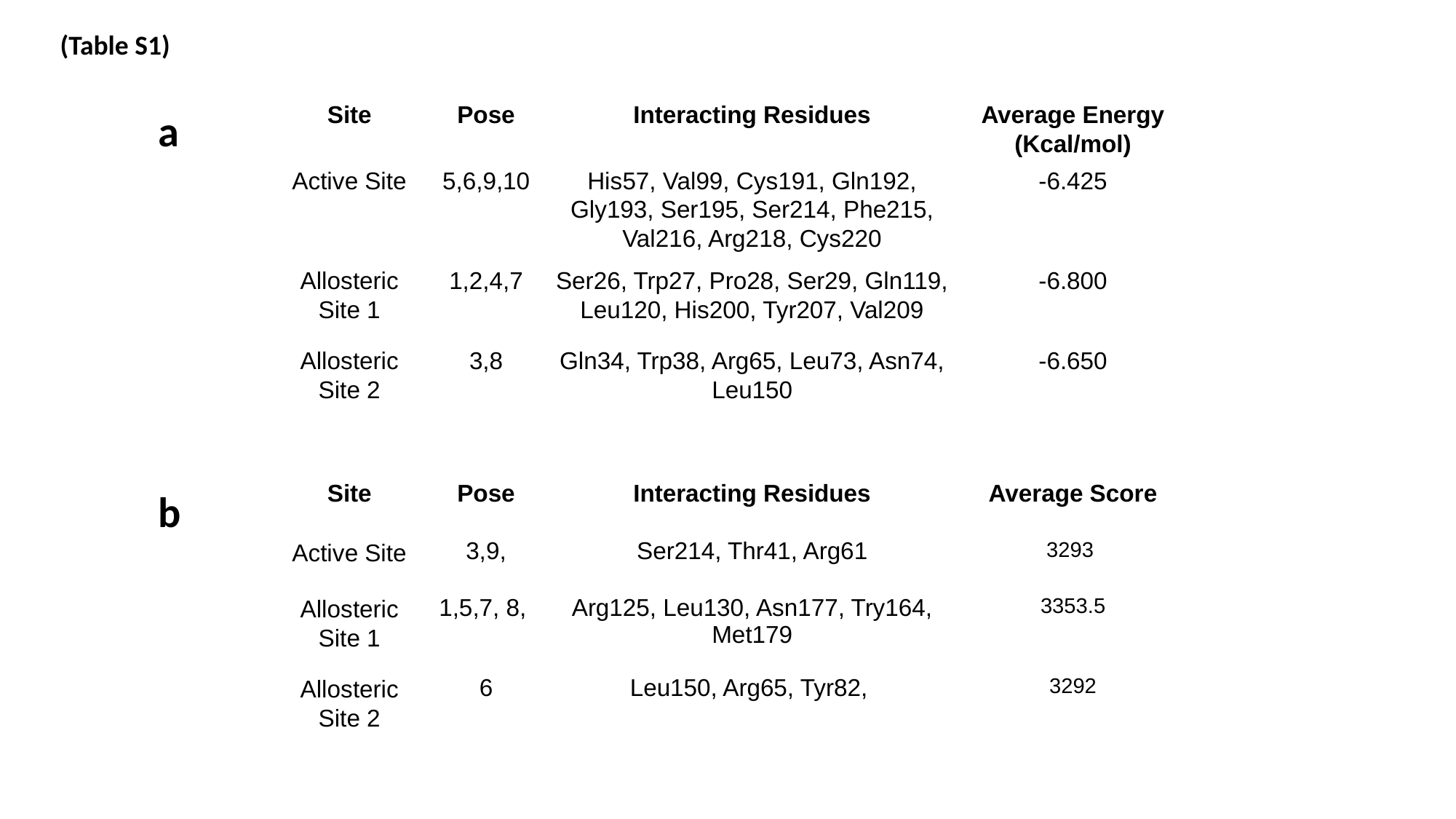

(Table S1)
a
| Site | Pose | Interacting Residues | Average Energy (Kcal/mol) |
| --- | --- | --- | --- |
| Active Site | 5,6,9,10 | His57, Val99, Cys191, Gln192, Gly193, Ser195, Ser214, Phe215, Val216, Arg218, Cys220 | -6.425 |
| Allosteric Site 1 | 1,2,4,7 | Ser26, Trp27, Pro28, Ser29, Gln119, Leu120, His200, Tyr207, Val209 | -6.800 |
| Allosteric Site 2 | 3,8 | Gln34, Trp38, Arg65, Leu73, Asn74, Leu150 | -6.650 |
b
| Site | Pose | Interacting Residues | Average Score |
| --- | --- | --- | --- |
| Active Site | 3,9, | Ser214, Thr41, Arg61 | 3293 |
| Allosteric Site 1 | 1,5,7, 8, | Arg125, Leu130, Asn177, Try164, Met179 | 3353.5 |
| Allosteric Site 2 | 6 | Leu150, Arg65, Tyr82, | 3292 |
